# Prefrontal-motor and somatosensory-motor cortical network interactions during reactive balance are associated with distinct aspects of balance behavior in older adults

**DOI:** 10.1101/2021.01.30.428951

**Authors:** Jacqueline A. Palmer, Aiden M. Payne, Lena H. Ting, Michael R. Borich

## Abstract

Heightened reliance on the cerebral cortex for postural stability with aging is well-known, yet the cortical dynamics of balance control, particularly in relationship to balance function, is unclear. Here we aimed to investigate motor cortical activity in relationship to the level of balance challenge presented during reactive balance recovery, and identify circuit-specific interactions between motor cortex and prefrontal or somatosensory regions to metrics of balance function that predict fall risk. Using electroencephalography, we assessed motor cortical beta power, and beta coherence during balance reactions to perturbations in older adults. We found that individuals with greater somatosensory-motor beta coherence at baseline and lower beta power evoked over motor regions following perturbations demonstrated higher general clinical balance function. At the group-level, beta coherence between prefrontal-motor regions reduced during balance reactions. Older adults with the highest post-perturbation prefrontal-motor coherence showed greater cognitive dual-task interference and elicited stepping reactions at lower perturbation magnitudes. Our results support motor cortical beta activity as a potential biomarker for individual level of balance challenge and implicate prefrontal-and somatosensory-motor cortical networks in different aspects of balance control in older adults. Cortical network activity during balance may provide a neural target for precision-medicine efforts aimed at fall-prevention with aging.

## Introduction

The development of balance impairment with aging is common but poorly understood. The neural mechanisms by which some individuals maintain high levels of activity while others suffer a debilitating loss of mobility and independence remain elusive. During the aging process there is a loss of automaticity in balance and mobility, where engagement of cortical resources for balance control interferes with the ability to perform cognitive and mobility tasks simultaneously (Jacobs & Horak, 2007; Lundin-Olsson et al., 1997; B. E. Maki & McIlroy, 2007). Previous studies have identified active cortical regions during continuous balance and walking tasks in older adults (Chang et al., 2016; Malcolm et al., 2020), but cortical oscillatory activity time-locked to destabilizing balance events, shown to be associated with balance ability in younger adults (Ghosn et al., 2020), or interactions between cortical regions reflecting information processing and integration during motor behavior have not been well investigated in older adults, who have increasing risk for falling with age. Such knowledge could identify effective neural control strategies during balance-correcting behavior in high-functioning older adults, and could be leveraged towards the development of precision medicine approaches for individualized fall prevention strategies among a heterogeneous older adult population. In the present study, we used electroencephalography (EEG) to measure time-locked cortical activity during standing balance reactions in a group of older adults across a range of individual balance abilities. We aimed to characterize the neural dynamics of cortical oscillatory activity and interactions between cortical regions during balance reactions, and test the relationship between individual cortical engagement strategy during balance reactions and distinct aspects of balance behavior that are predictive of falls in older adults.

Balance control is multifactorial. A variety of tests measure different aspects of balance control, whose neural underpinnings are not well understood. Growing evidence shows that engagement of cortical resources during balance is an indicator of fall risk in older adults, where a concurrent cognitive task shifts cortical resources away from balance control (Lundin-Olsson et al., 1997; Montero-Odasso et al., 2012; Shumway-Cook et al., 1997; Woollacott & Shumway-Cook, 2002). Commonly used clinical tests, such as the miniBEST, that assess a myriad of aspects of postural control including single-task, cognitive dual-task, and reactive balance, have high clinical utility, but are nonspecific, lack precision, and can impose a ceiling effect on individuals with higher balance ability (Marques et al., 2016). The ability to react to a loss of balance is a key factor that ultimately determines whether an individual will sustain a fall. These balance recovery mechanisms can be profoundly impaired in older adult populations (for comprehensive review see (B. E. Maki & McIlroy, 1996; Brian E. Maki & McIlroy, 2006)) and are strongly linked to fall risk (Chandler et al., 1990; Wolfson et al., 1986). The link between balance recovery ability and fall risk has prompted researchers to quantify reactive balance capacity as the initiation of stepping responses during postural destabilization of a given magnitude, where individuals with greater reactive balance impairment require stepping reactions at lower levels of balance perturbations (Jensen et al., 2001; Mille et al., 2003). Reactive balance recovery also provides a unique paradigm to assess cortical activity dynamics that are time-locked to balance behavior using electroencephalography (EEG). In the present study, we aimed to quantify individual reactive balance capacity by increasing the magnitude of balance perturbations to the point where balance challenge exceeded the capacity for an individual to produce feet-in-place reactions, necessitating a later-phase reactive stepping response that is likely cortically-mediated (B. E. Maki & McIlroy, 2007). Previously, our lab used EEG to assess cortical activity during balance reactions in younger adults and found larger evoked cortical responses in individuals with lower balance performance on a beam walking task (Ghosn et al., 2020; Payne & Ting, 2020). Further, evoked cortical activity was dissociable from evoked muscle activity, suggesting a potential cortical motor contribution to later-phase reactive balance control (Payne et al., 2019). However, it is unclear whether older adults engage similar cortical strategies during balance reactions, and whether behavioral assessments of fall risk used in older adult populations assess similar or distinct aspect of balance control. Moreover, understanding the neural control strategies underpinning of the various aspects of balance would be useful for the development of fall prevention treatments within a precision-medicine framework.

Investigating the information processing between motor cortical and other brain regions during whole-body balance-correcting behavior could provide valuable information about the time course of circuit-specific cortical contributions to balance control that cannot be observed using measures of cortical activity during rest or static balance alone. Neural oscillations in the beta frequency band (13-30Hz) are a prominent feature of motor behavior (van Wijk et al., 2012; Zaepffel et al., 2013) (Engel & Fries, 2010). In older adults, and in age-related neurodegenerative disease such as Parkinson’s disease, abnormal movement-related beta oscillatory modulation has been associated with slowed and impaired volitional motor activity (P. Brown, 2007; Johari & Behroozmand, 2020). Consistent with age-related increases in cortical recruitment during motor tasks (R. D. Seidler et al., 2010), older adults had greater movement-related modulation of beta activity compared to younger adults during volitional manual movements (Rossiter et al., 2014). Age-related differences in functional connectivity between cortical regions may, in part, explain differences in movement-related cortical beta activity in older adults, which has been observed in older individuals with Parkinson’s disease and stroke (Grefkes et al., 2008; Rowe et al., 2002). Functional connectivity analyses performed in during resting motor states suggest that the role of neural interactions between cortical regions may be circuit-specific (Langan et al., 2010; R. Seidler et al., 2015; Solesio-Jofre et al., 2014). However, the functional role of cortical beta oscillations and circuit-specific functional connectivity in the aging brain remains less clear in behavioral contexts, particularly whole-body balance reactions.

Heightened cortical activity in somatosensory and motor regions has been observed in older adults compared to younger individuals and associated with upper limb motor function, but it is unclear whether sensorimotor cortical activity is also associated with balance function. Centrally-mediated sensorimotor processing within the cortex may preserve balance control in the presence of age-related loss of somatosensory function (Zhang et al., 2011) and the loss of automaticity of balance and mobility behavior via subcortical mechanisms (Clark, 2015). Older adults consistently show a greater extent of activation in somatosensory and motor cortical brain regions during a myriad of single-segment limb motor tasks or tasks that mimic whole-body task performance (e.g., virtual reality or mental imagery) compared to younger adults (Cassady et al., 2020; Goble et al., 2012; Heuninckx et al., 2008; Mattay et al., 2002; Zwergal et al., 2012). Increases in somatosensory and motor cortical activity with aging have also been accompanied by greater functional connectivity between somatosensory and motor regions during finger tapping tasks, suggesting causal network interactions between these brain regions (Cassady et al., 2020). Though cortical interactions between somatosensory and motor regions have been historically challenging to study in the context of whole-body behaviors, higher levels of sensorimotor cortical activity have been positively associated with interlimb coordination performance (Heuninckx et al., 2008), potentially implicating a beneficial functional role for sensorimotor cortical control in bilateral limb motor performance during standing balance and mobility. In this study, we tested this theory by measuring somatosensory-motor interactions during reactive balance and its association with balance ability in older adults.

Prefrontal cortical brain regions subserving cognitive executive function and working memory appear to play an increased functional role in balance control with aging, and may limit simultaneous cognitive/balance task performance and the upper limit of individual motor performance capacity in older adults. Cognitive interference during balance and mobility in older adults (L. A. Brown et al., 1999; Leone et al., 2017; Morris et al., 2016; Rankin et al., 2000; Woollacott & Shumway-Cook, 2002) suggests the development of overlapping cortical control mechanisms for cognitive and motor control processes with aging (Cid-Fernández et al., 2014; Ren et al., 2013), implicating an increased role of the prefrontal cortex for balance control. Older adults show greater prefrontal cortical activity during a wide range of motor tasks, including interlimb coordination tasks (Heuninckx et al., 2008) and steady-state walking (Chen et al., 2017; Hawkins et al., 2018; Mirelman et al., 2017). Whether age-related differences in prefrontal cortical activity play a beneficial (i.e. compensatory) (Clark et al., 2019) or detrimental (i.e. age-related neural dedifferentiation) (Gagnon et al., 2019; Payer et al., 2006) role in motor function in older adults remains controversial (for review see R. D. Seidler et al., 2010) and may depend on the context and challenge of the motor task (e.g. level of complexity and difficulty) (Clark, 2015). In contrast to younger adults, older adults utilize motor control strategies that require higher levels of cognitive processing, which are effective at slower speeds but less effective during fast speed motor performance (Boisgontier & Nougier, 2013). Additionally, individuals who show greater prefrontal cortex activity at low levels of task difficulty appear to have limited ability for additional prefrontal resource recruitment as the complexity and challenge of the task increases, ultimately limiting the upper end of performance capacity (Hawkins 2018). In this study, we tested whether the level of engagement of prefrontal-motor cortical networks during balance reactions at a matched perturbation magnitude across participants was associated with the level of cognitive dual-task interference and reactive balance performance under the most challenging perturbation conditions.

In the present study we aimed to investigate motor cortical beta activity and circuit-specific interactions between motor cortex and prefrontal or somatosensory brain regions over the time course of standing balance recovery in older adults. We tested the relationship between motor cortical beta power and beta coherence of somatosensory-motor and prefrontal-motor networks with three key aspects of functional balance ability in older adults, namely general clinical balance function, cognitive dual-task interference, and upper-end reactive balance capacity measured as reactive step threshold. We hypothesized that somatosensory-motor cortical networks would contribute to balance control in older adults and therefore greater somatosensory-motor coherence would be present in older adults with higher balance ability. We predicted that individuals with greater recruitment of prefrontal-motor circuits at a matched perturbation magnitude would be more susceptible to cognitive dual task interference, showing more slowing in their mobility performance during simultaneous cognitive task performance, and have lower reactive balance capacity, measured as eliciting of a stepping reaction at lower perturbations as magnitude was increased.

## Methods

### Participants

Sixteen individuals were recruited from the local Atlanta community to participate in this study. All participants completed a single testing session consisting of clinical balance assessments and neurophysiologic testing during a standing balance perturbation series. Inclusion criteria included above the age of 50, the ability to walk at least 10 meters without the assistance of another person, the ability to stand unassisted for at least 3 minutes, and the cognitive ability for informed consent. No participants in the present study used an assistive device for ambulation. Participants were excluded if they had the diagnosis of any neurologic condition, any musculoskeletal condition that affected their standing or walking, peripheral neuropathy, or pain affecting standing or walking. The experimental protocol was approved by the Emory University Institutional Review Board and all participants provided written informed consent.

### Behavioral balance assessments

#### General clinical balance function

Upon arrival to the lab and prior to neurophysiologic instrumentation, participants completed the miniBEST to assess general clinical balance function. The miniBEST is a validated and commonly used clinical assessment for assessing static and dynamic balance ability and fall risk in elderly adults (Marques et al., 2016). Briefly, the miniBEST assesses domains of anticipatory balance control, reactive postural control, sensory orientation, and dynamic gait using a likert subscale of 0-2 for each domain, where higher scores indicate better performance. The total sum of the itemized subscale scores represented miniBEST total score, with a maximum possible score of 28 (taking the lower of two scores for items scored separately for left and right legs).

#### Dual-task interference

Dual-task interference was assessed during a clinical Timed-Up-and-Go (TUG) test, a validated clinical test for fall risk assessment in elderly populations (Tang et al., 2015). Participants started in a seated position with their back against the back of a chair. When the clinician verbally cued a “Go” signal, the participant stood up from the chair, walked 3 meters until both feet crossed over a taped line on the floor and walked back to the chair, with the time stopping when their back came in contact with the back of the chair. Next, the participants were instructed to repeat the TUG test while performing the secondary cognitive task of verbally counting backwards by 3’s starting at a random integer number between 20 and 100 verbally stated by the experimenter immediately following the “Go” signal. Participants were instructed that this was a timed test and to “walk as fast as you safely can” during both single and dual-task performance. Dual-task interference (DTI) was quantified as (Plummer & Eskes, 2015):

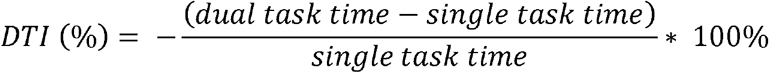

Negative DTI values indicate slower TUG performance during dual task relative to the single task condition. All clinical testing was administered by a licensed physical therapist.

#### Behavioral reactive balance capacity

We assessed individual behavioral reactive balance capacity by determining the lowest perturbation magnitude which elicited unintentional stepping reactions in approximately 50% of the trials, defined as the step threshold. The step threshold testing was performed after a seated rest break following the first series of perturbations described above to avoid initial behavioral adaptation effects that could occur at the start of the moving platform series. Participants stood on the platform with the same instructions to attempt to respond with a feet-in-place strategy and arms crossed in place at the chest. Forward direction perturbations were delivered starting at a magnitude of 8 cm with a jittered inter trial interval of 15-60 seconds. To reduce anticipation of perturbation direction, backwards directional perturbations were also randomly administered with this perturbation series. If the participant successfully completed 3 consecutive feet-in-place trials, then the perturbation magnitude was scaled up by 1cm displacement, with proportional increases in perturbation velocity and acceleration. This procedure was repeated, scaling up the perturbation magnitude until a step reaction occurred. At that point, the perturbation magnitude was held constant at this level for the next 10 perturbations. If the participant elicited 5 step reactions in a row at this perturbation magnitude, the perturbation magnitude was scaled down by 0.5 magnitude level. The step threshold was defined as the perturbation magnitude at which the participant utilized an unintentional step response strategy in approximately 5 out of 10 (50%) of trials. During this perturbation series, the experimenter closely monitored the participant’s real-time force data during baseline quiet standing to ensure the same baseline standing position. If the participant attempted to adjust their posture in anticipation of a perturbation (e.g. increase stance width or forward trunk lean), the experimenter cued the participant to return to their normal baseline standing posture prior to perturbation delivery.

### Balance perturbations

#### Matched perturbation level protocol

Participants stood with bare feet in the middle of a moveable custom platform (Factory Automation Systems, Atlanta, GA) while support-surface translational perturbations were delivered in an unpredictable direction and at unpredictable timing. Twenty-four perturbations of equal magnitude (7.5 cm, 16.0 cm/s, 0.12 g) were delivered in the forwards direction to elicit a backwards center of mass displacement relative to the base of support. The scaling of these perturbation parameters was selected to ensure that platform deceleration did not occur until 500 ms after perturbation onset to minimize changes in cortical and motor output originating from a deceleration response (Ghosn et al., 2020; McIlroy & Maki, 1994). Because we aimed for each participant to sustain identical perturbations, we selected this lower-level perturbation magnitude as a level of postural destabilization that could be successfully completed by most older adults using a feet-in-place strategy (Figure 1). To reduce the directional anticipation and time of perturbation onset, we included three additional perturbation directions (backwards, 45 degrees right posterolateral, 45 degrees left posterolateral) using the same scaled parameters into the perturbation series in a pseudorandomized order, where a perturbation of the same magnitude had no more than 2 consecutive occurrences. To reduce anticipation of precise perturbation onset, perturbations were delivered at a jittered intertrial interval, with 15 – 60 seconds between each perturbation onset. Real-time EEG activity and force feedback was also monitored by the experimenter to ensure that the participant returned to baseline levels of cortical and muscle activity and maintained the same baseline body position. Participants were asked to attempt to recover balance while maintaining both feet in place and arms crossed at their chest. If a participant executed an unintentional stepping reaction in response to a perturbation by visual determination by the experimenter in real time, the trial was marked for offline confirmation and exclusion based on ground reaction forces. Participants took a seated rest break every 8 minutes during balance perturbation testing, or more frequently if the participant requested a break or reported or showed signs of fatigue during testing.

**Figure 1.**
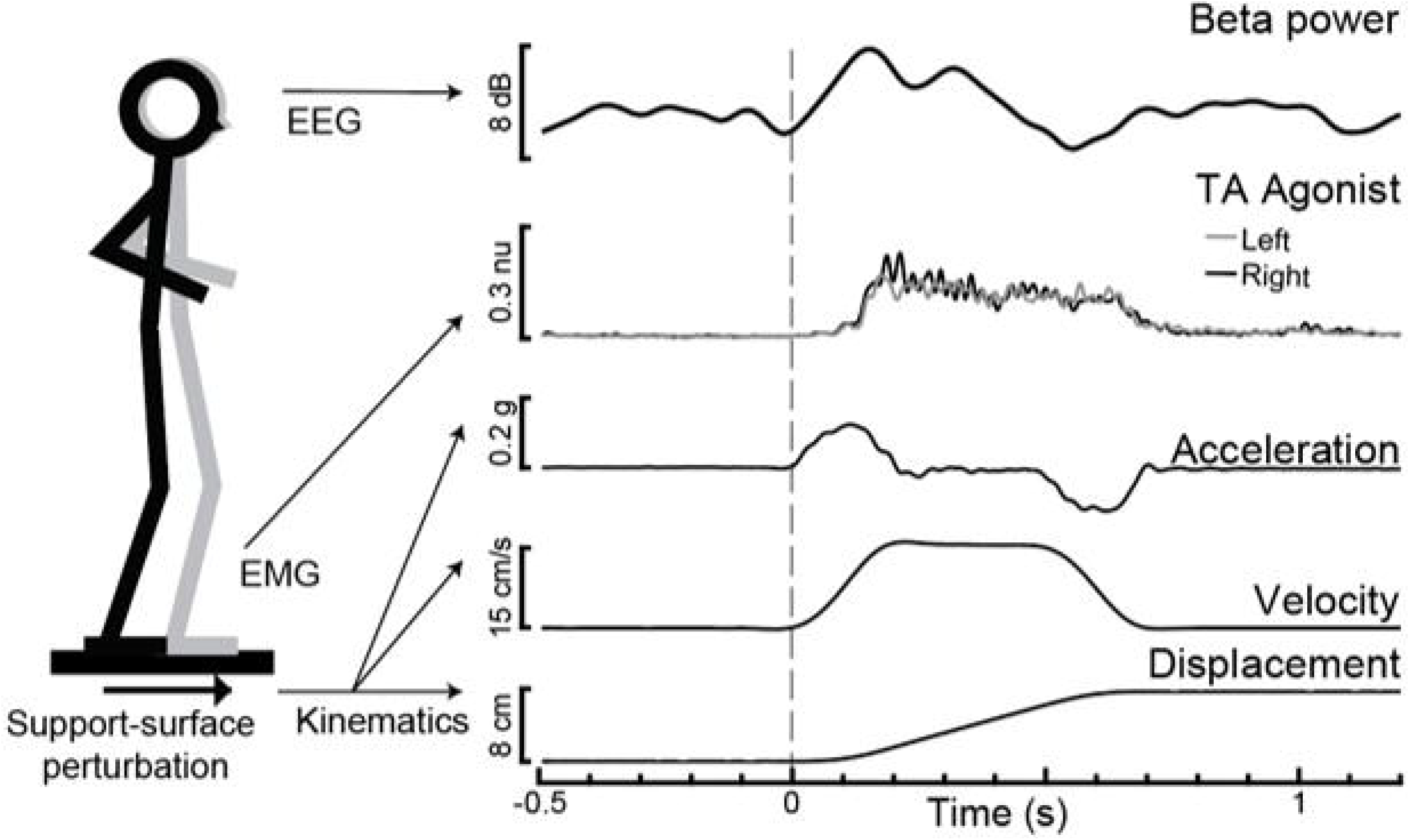
Experimental paradigm with evoked motor cortical beta power (Cz) and tibialis anterior (TA) agonist muscle activity with support-surface perturbation kinematics for an exemplar participant.

### Electroencephalography (EEG) data collection and analyses

During the matched-level balance perturbations, EEG signals using a 64-channel active electrode cap (ActiCap, Brain Products GmbH, Gilching, Germany) connected to an ActiCHamp amplifier (Brain Products, GmbH). Data were continuously recorded and online referenced to the FCz channel (Recorder, Brain Products, GmbH). EEG signals were digitized with a 24-bit analog-to-digital converter and an online 20 kHz low-pass filter and were sampled at 1000Hz and stored for offline analyses.

#### Preprocessing

All EEG data were preprocessed using freely available functions from the EEGlab toolbox (Delorme & Makeig, 2004). The time-locked continuous data were imported into EEGlab and filtered with high-pass cutoff of 1Hz and low-pass cutoff of 100Hz. Next, the events with trigger labels for successful trials (no step) in the forward direction of platform translation were selected (−2 to 3seconds relative to the time of the platform movement onset trigger at t=0) and any trial that was contaminated by artifacts was removed from analysis. Highly contaminated channels outside of the channels used for primary analyses (Cz, CPz, AFz) were identified by visual inspection and removed from the recordings. On average, 62 of 65 channels remained for analyses (SD ± 3.4); range 53-65). The removed electrodes were then interpolated using the *pop_interp* function in EEGlab. The *Cleanline* plugin for EEGLAB was applied to the continuous data to remove line noise (60 Hz). Next, the data were epoched (−1 to 2 seconds relative to perturbation onset). We then applied Independent Component Analysis (ICA) to remove non-neural artifacts of the EEG signal (e.g. muscle activity, motion, eyeblinks). Using as few as 32 electrodes, ICA has been shown to effectively disentangle motor, sensory, and cognitive processes, even when they are occurring simultaneously with overlapping scalp distributions and frequency properties (Makeig et al., 2004). Non-neural artifacts (e.g. muscle activity, motion artifact, eye blinks, cardiac rhythm) were identified using an automatic component selection TESA algorithm (Rogasch et al., 2017) and visually confirmed for accuracy. The remaining components were retained for subsequent analyses.

#### Quantification of cortical beta power

Time-frequency decomposition analyses were used to quantify changes in beta oscillatory power in response to balance perturbations for the vertex electrode overlying the primary motor cortex (Cz). We used the wavelet time-frequency analyses function (pop_newtimef.m) in EEGLAB to quantify beta oscillatory power across all forward perturbations within each participant. We used a sliding window of 256ms to measure power at each frequency using a tapered Morlet wavelet. The lowest frequency (12Hz) used three oscillatory cycles, which increased up to six cycles used at the highest frequency (50Hz). The event-related spectral perturbation (ERSP) quantified oscillatory power (Makeig, 1993) at 10 linearly spaced frequencies (12Hz to 50Hz) at 14ms intervals throughout the perturbation trials. To index power in the beta frequency domain, the mean of the ERSP values across four sampled frequencies centered on 16Hz, 20Hz, 24Hz, and 29Hz were computed for each participant. We then computed the mean baseline beta power (−500 to 0ms before perturbation onset) and the peak of the perturbation-evoked change in beta power (100-500ms) relative to baseline for each participant. Based on our previous study which found differences between early and later portions of motor cortical beta activity responses as a function of balance ability in younger adults (Ghosn et al., 2020), we further identified peak perturbation-evoked beta power within an early (100-300ms) and later (300-500ms) time window of the response.

#### Quantification of circuit-specific cortical coherence

Imaginary part of coherence (IPC) analyses (Nolte et al., 2004) were used to quantify the phase-shifted synchrony of oscillatory activity within the beta frequency range (13-30Hz) between two pairs of vertex electrodes approximately overlying lower limb regions of the primary motor cortex (Cz) with 1) primary somatosensory (CPz) and 2) dorsolateral prefrontal cortical (AFz) regions. Given the close anatomical proximity of our brain regions of interest, we chose this methodologically conservative coherence analysis approach to minimize the risk of artificially inflated cortical coherence due to volume conduction by requiring a phase lag between distinct source signals (Nolte et al., 2004). Analyses were performed using custom routines in MATLAB, using an upper end frequency cut-off of 50 Hz, segment length of 768, and overlay of 0.9 (Palmer et al., 2020) to yield a frequency domain resolution of 1.3 Hz and time domain resolution of 76.8 ms. The pre-perturbation IPC value within the 400 ms prior to perturbation onset (−400 to 0 ms) was computed from the mean of the six segments prior to t=0, with the first segment centered at −388ms and the last segment at −8ms. The time window of 100-500 ms post-perturbation onset was selected because it captured the earliest occurrence of motor cortical beta power change within the group (Figure 2) and would not be affected by the platform deceleration after 500 ms (Figure 1). The post-perturbation (100-500 ms) IPC value was computed from the mean of the first five segments relative to t=0, with the first segment centered at 144ms and the last segment at 448ms.

**Figure 2.**
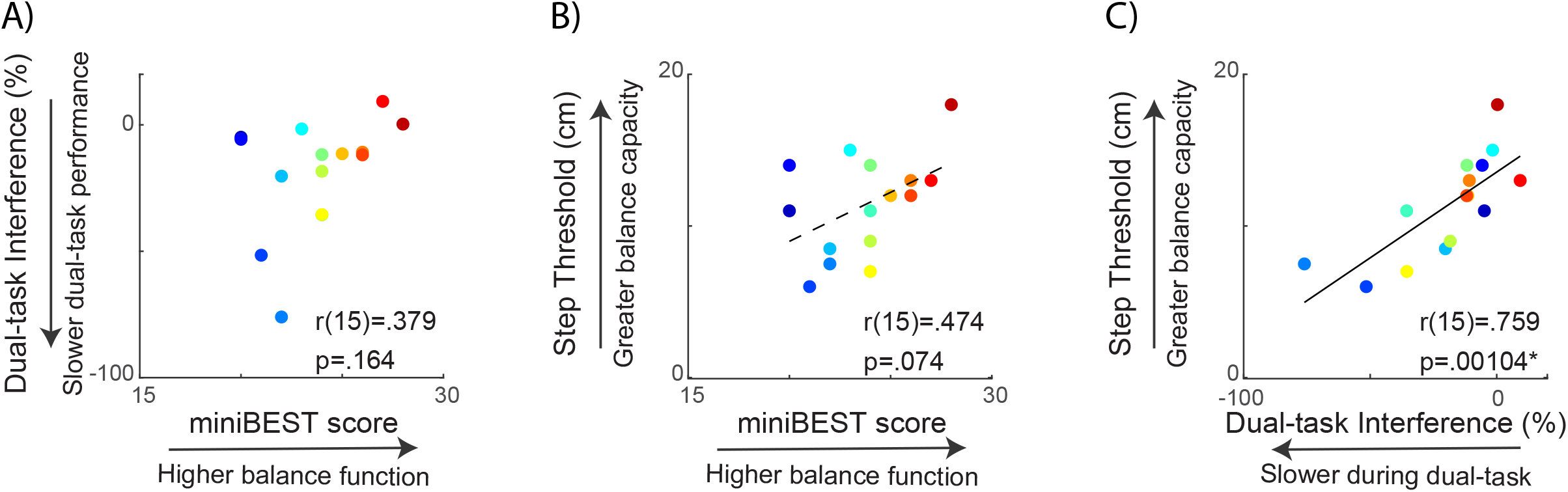
Relationship between clinical behavioral balance function and reactive step threshold. MiniBEST score was not significantly associated with cognitive dual-task interference **(A)** or reactive step threshold **(B).** Cognitive dual task interference was positively correlated with reactive step threshold, where older adults with more slowing during dual task performance had lower reactive step thresholds **(C)**.

### Statistical analyses

We used Kolmogorov-Smirnov and Levene’s tests to test for normality and homogeneity of variance in balance behavior, cortical beta power, and cortical beta coherence data. We tested the relationship between clinical balance behavioral measures of miniBEST, dual task interference, and reactive step threshold using Pearson product-moment correlation coefficients. We tested the modulation of motor cortical beta power, somatosensory-motor beta coherence, and prefrontal-motor beta coherence between pre-and post-perturbation time windows during balance reactions using paired t-tests. We tested the relationship between perturbation-evoked motor cortical beta power and miniBEST score using Pearson product-moment correlation coefficients. We used Pearson product-moment correlation coefficients to test relationships between pre-and post-perturbation S1-M1 and PF-M1 beta coherence versus miniBEST, dual task interference, and step threshold. Statistical analyses were performed using Statistical Package for Social Sciences version 27 (IBM Corp, Armonk, NY) with an *a priori α* level set to .05.

## RESULTS

Fifteen (age: 69 ± 8 years, 11 female, Table 1) out of 16 participants were able to complete the protocol. One participant withdrew from the study due to high levels of anxiety and fear of falling during platform movement and was excluded from all analyses. One participant was only able to successfully execute 4 trials in the forward direction without a stepping reaction; for this participant, successful no step trials in the backward direction were included in EEG analysis to standardize the number of trials across participants in EEG data analysis. One other participant was unable to successfully respond to balance perturbations at the 7.5 cm magnitude with feet in place in any direction; for this participant, the perturbation parameters were scaled down to 7 cm magnitude displacement (7 cm, 15 cm/s, 0.12 g), a level where feet-in-place trials could be successfully executed. As a group, participants were able to successfully recover balance without stepping in 91.3±1.4% of the trials on average. Thus, an average of 21.9±5.0 trials were used for EEG data analyses across participants. The EEG recordings of one other participant had excessively high impedances (>50 kOhm) secondary to use of an oil-based hair product on the scalp prior to testing; these data were excluded from EEG analyses but retained for balance behavioral analyses.

**Table 1.**
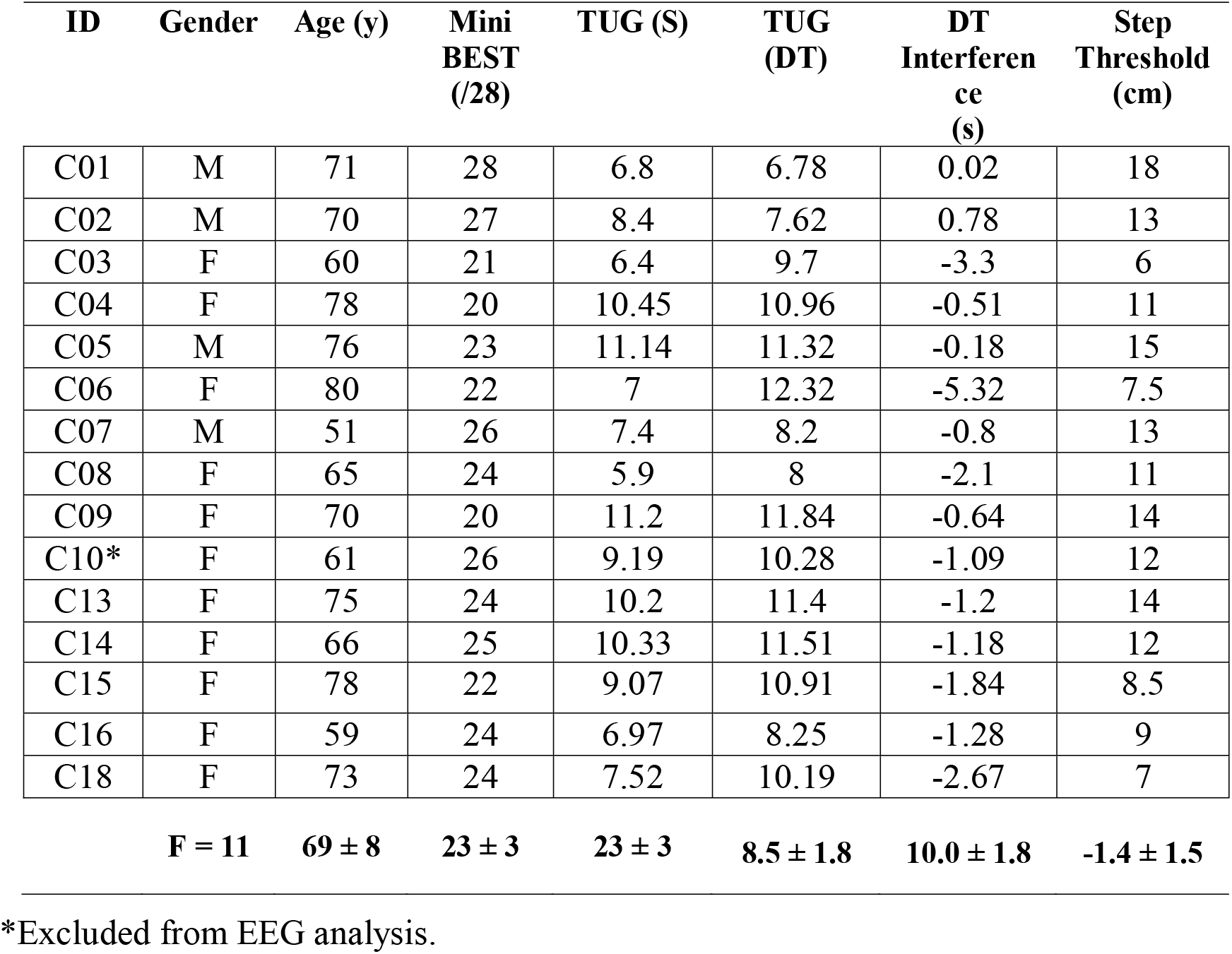
Participant characteristics and balance behavior

### Assessment of clinical balance function and reactive balance capacity

When testing the relationship between behavioral balance measures, we found a strong relationship between cognitive dual-task interference and reactive step threshold, while there was no significant relationship between general clinical balance function, measured as the miniBEST, and either cognitive dual-task interference or reactive step threshold. Individual miniBEST scores were not associated with cognitive dual-task interference (*r*(15)=.379, *p*=.164) (**Figure 2A**) or reactive step thresholds (*r*(15)=.474, *p*=.074) or (**Figure 2B**). Individual cognitive dualtask interference was positively associated with reactive step threshold (*r*(15)=.759, *p*=.001), where individuals with greater slowing during the cognitive dual-task condition elicited reactive stepping responses at lower perturbation magnitudes (**Figure 2C**).

### Perturbation-evoked motor cortical beta power

We found that older adult with greater later-phase perturbation-evoked motor cortical beta power had higher levels of general clinical balance function, as measured by the miniBEST. Balance perturbations elicited an increase in motor cortical beta oscillatory power from the baseline mean (27 ± 3 dB) to a post-perturbation (100-500 ms) mean (36 ± 7 dB) (*t*=-5.64, *p*<.0001) (**Figure 3**). There was no relationship between perturbation-evoked beta power during the overall (100-500 ms) (*r*(14)=.49, *p*=.074) or early-phase (100-300ms) (*r*(14)=.43, *p*=.13) time window (**Figure 4**). Later-phase perturbation-evoked beta power was inversely associated with miniBEST (*r*(14)=.59, *p*=.026), where individuals with higher perturbation-evoked beta power during the later-phase of balance reactions had lower clinical balance function (**Figure 4)**. We did not observe an association between perturbation-evoked beta power during any time window and cognitive dual task interference or reactive step threshold (p>.05).

**Figure 3.**
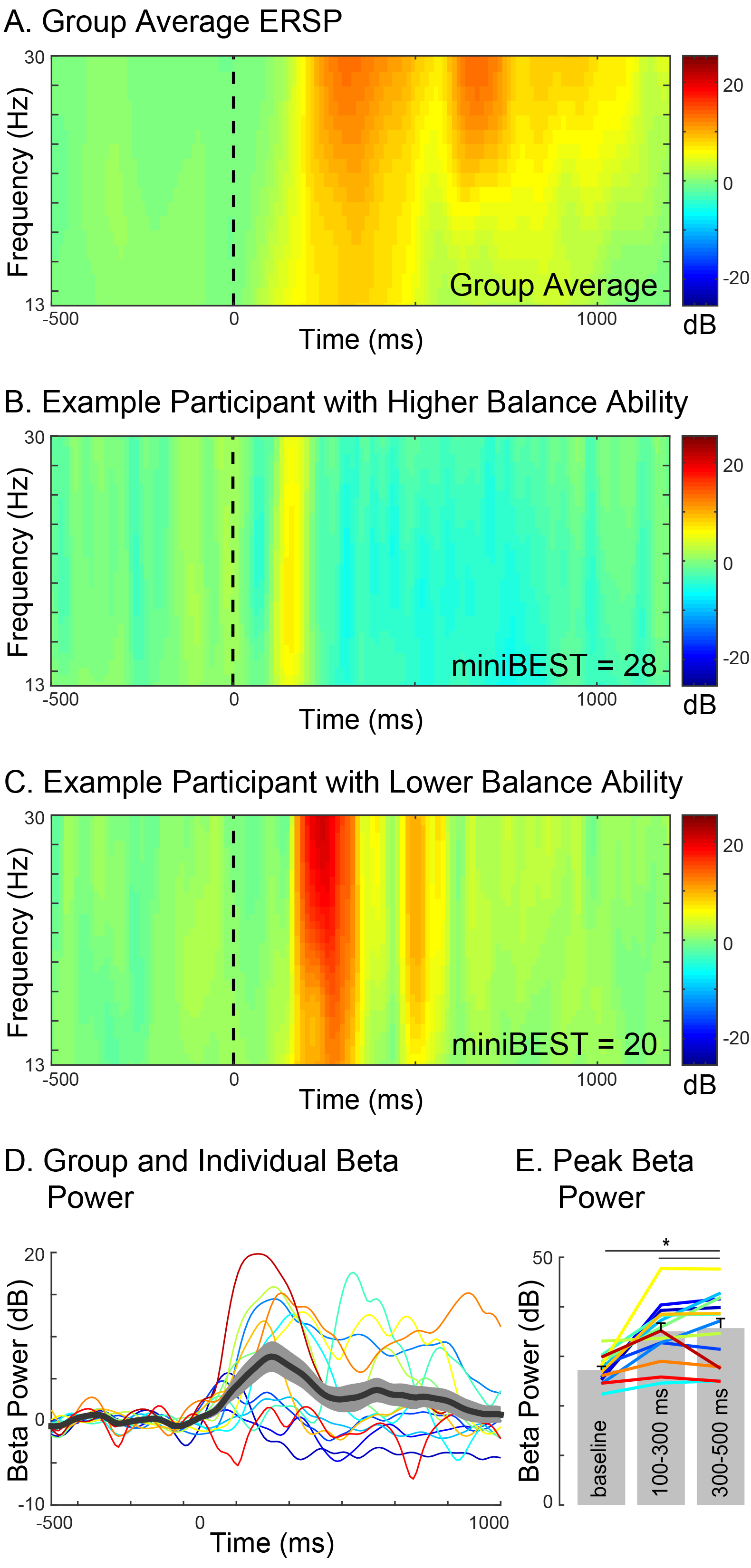
Motor cortical beta oscillatory power (Cz) during reactive balance responses. Group level event-related spectral perturbation (ERSP) in the beta frequency range (13-30Hz) **(A)** ERSP in two exemplary individuals with higher (**B**) and lower (**C**) miniBEST score. Time course of beta power response across individuals **(D).** Beta power increased from pre-to postperturbation during both 100-300ms and 300-500ms time bins(*p<.001) **(E).**

**Figure 4.**
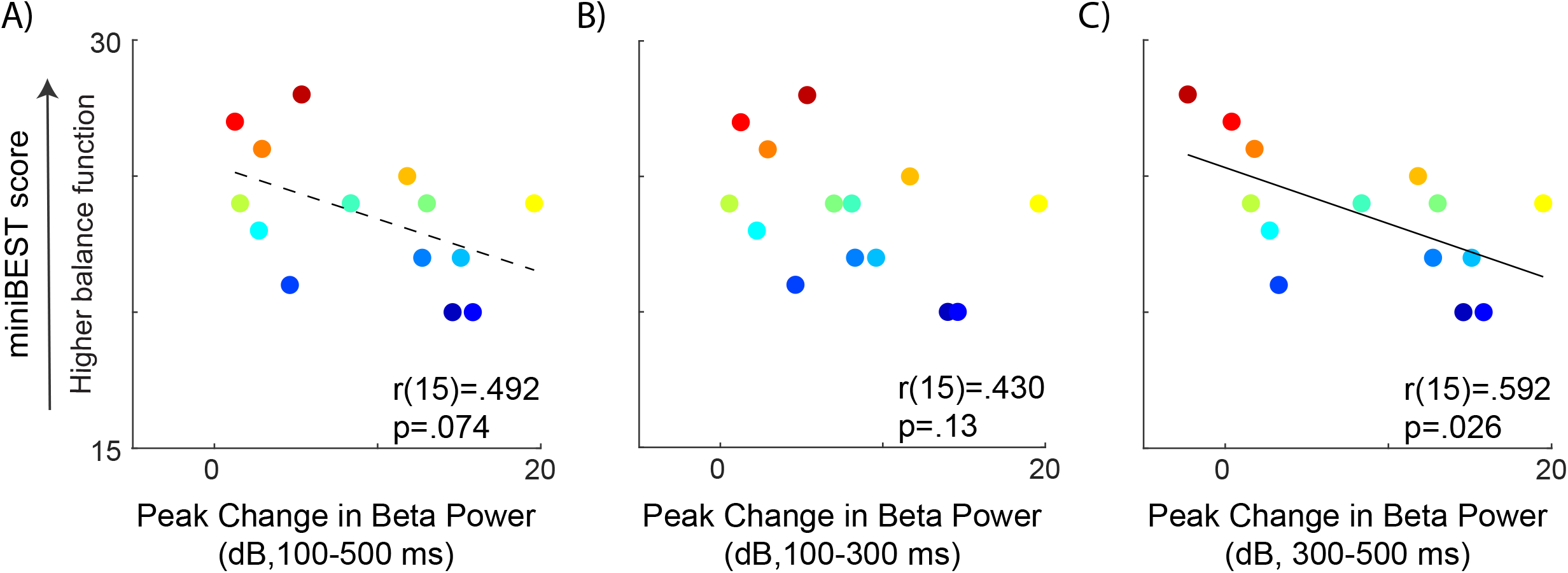
Relationships between perturbation-evoked beta power (baseline subtracted) during overall (**A**), early (**B**) and later-phase balance reactions (**C**). No relationships were observed between perturbation-evoked beta power during overall (100-500ms) or early-phase (100-300ms), but later-phase (300-500ms) perturbation-evoked beta power was negatively associated with miniBEST score. Individual color bar scaled by miniBEST score throughout (red=highest miniBEST, blue = lowest miniBEST).

### Perturbation-evoked motor cortical coherence

Older adults who showed greater somatosensory-motor coherence had higher miniBEST scores, while older adults who showed greater prefrontal-motor coherence had greater levels of cognitive dual-task interference. Though balance perturbations did not elicit a change in somatosensory-motor beta coherence at the group level (*t*=1.63, *p*=.13), responses were highly variable between individuals, particularly between those with higher versus lower miniBEST scores (**Figure 5A**). Group-level prefrontal-motor beta coherence reduced post-perturbation (*t*=2.38, *p*=.03), but this response was also variable between individuals, particularly between those with less versus more cognitive dual-task interference (**Figure 5B**).

**Figure 5.**
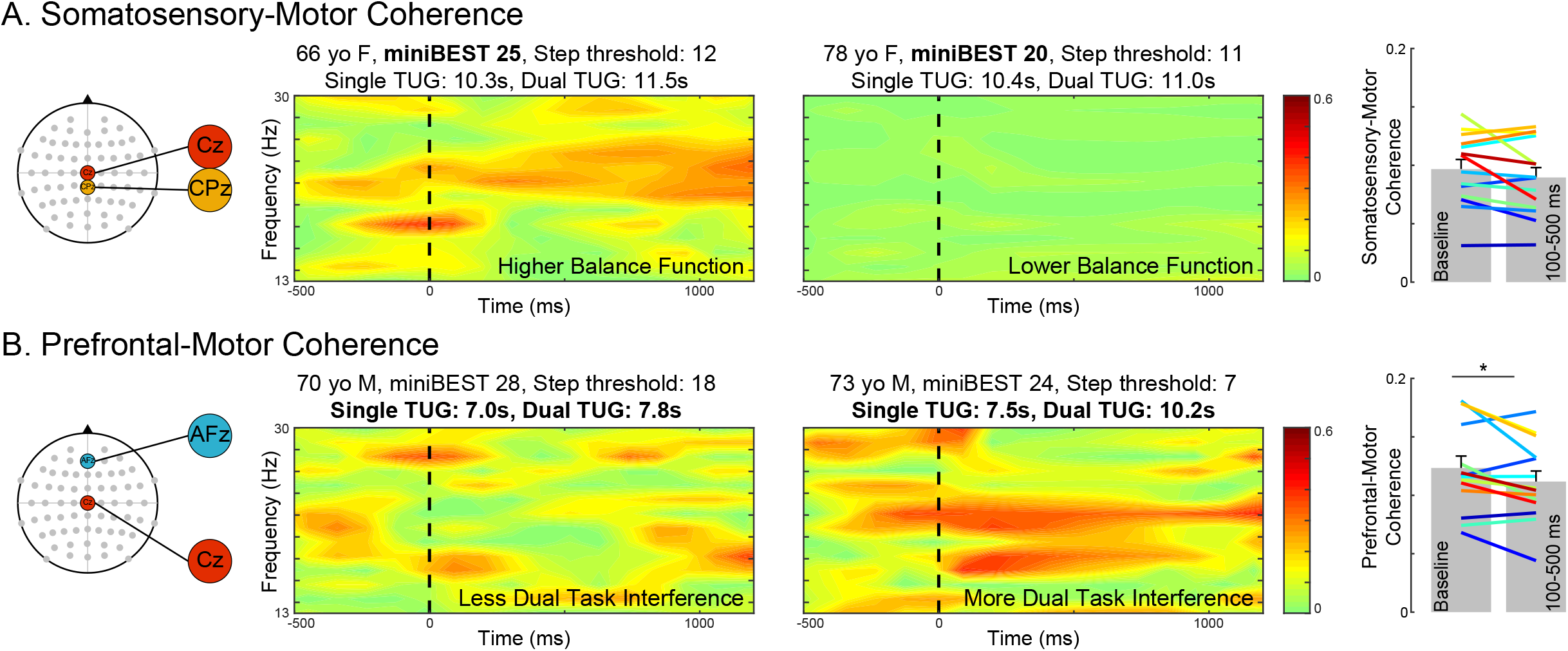
Somatosensory-motor and prefrontal-motor cortical beta coherence during balance reactions. Somatosensory-motor beta coherence was different between individuals with high versus low miniBEST score, and did not change following balance perturbation **(A).** Prefrontal-motor beta coherence was different between individuals with less versus more cognitive dualtask interference, and reduced post-perturbation (100-500ms) (\**p*=.03) **(B).** Broken line indicates onset of perturbation.

We observed a positive relationship between somatosensory-motor beta coherence and miniBEST score pre-perturbation (*r*(14)=.637, *p*=.014) but not post-perturbation (*r*(14)=.51, *p*=.062), such that older adults with greater baseline somatosensory-motor coherence had higher clinical balance function (**Figure 6A**). There was no relationship between somatosensory-motor coherence and step threshold at any time point (pre-perturbation, *r*(14)=.056, *p*=.849; postperturbation, *r*(14)=.034, *p*=.907) (**Figure 6B)** or somatosensory-motor coherence and cognitive dual-task interference at any time point (pre-perturbation, *r*(14)=.202, *p*=.489; post-perturbation, *r*(14)=.072, *p*=.806) (**Figure 6C**).

**Figure 6.**
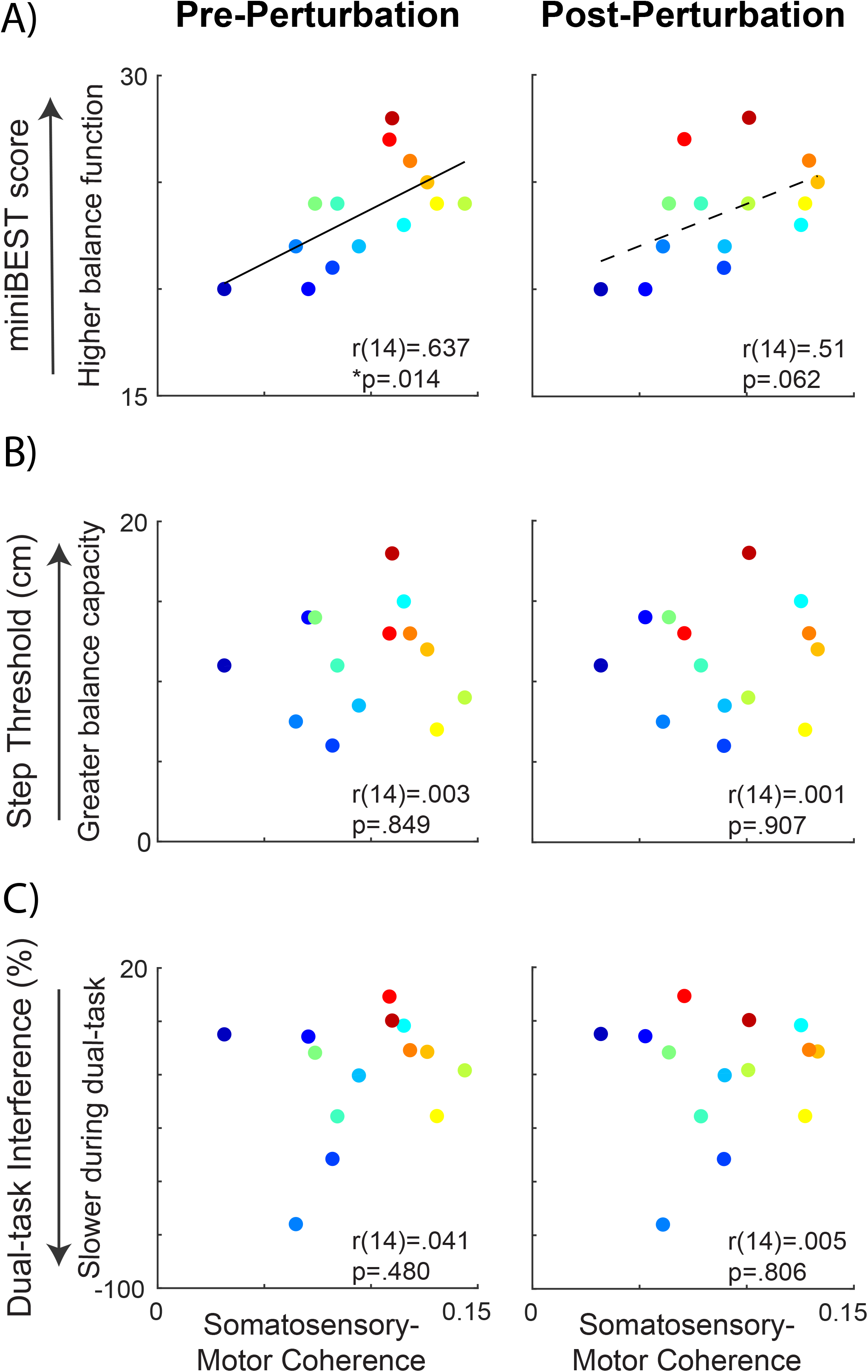
Somatosensory-motor cortical beta coherence pre-and post-perturbation and relationships to balance behavior. Somatosensory-motor coherence was positively associated with miniBEST at pre-perturbation (\**p*=.014) but not post-perturbation time-points **(A).** Somatosensory-motor coherence was not associated with reactive step threshold **(B)** or cognitive dual task interference **(C)**.

While there was no relationship between prefrontal-motor coherence and miniBEST (**Figure 7A**), we observed a negative relationship between post-perturbation prefrontal-motor coherence and both reactive step threshold (*r*(14)=-.543, *p*=.045) (**Figure 7B**) and cognitive dual task interference (*r*(14)=-.600, *p*=.025) (**Figure 7C**), where older adults with greater prefrontal-motor beta coherence elicited stepping reactions at smaller perturbation magnitudes and showed greater performance decline cognitive loading. This relationship was not present at the pre-perturbation time point (dual-task interference (*r*(14)=-.325, *p*=.257); step threshold (*r*(14)=-.390, *p*=.168)).

**Figure 7.**
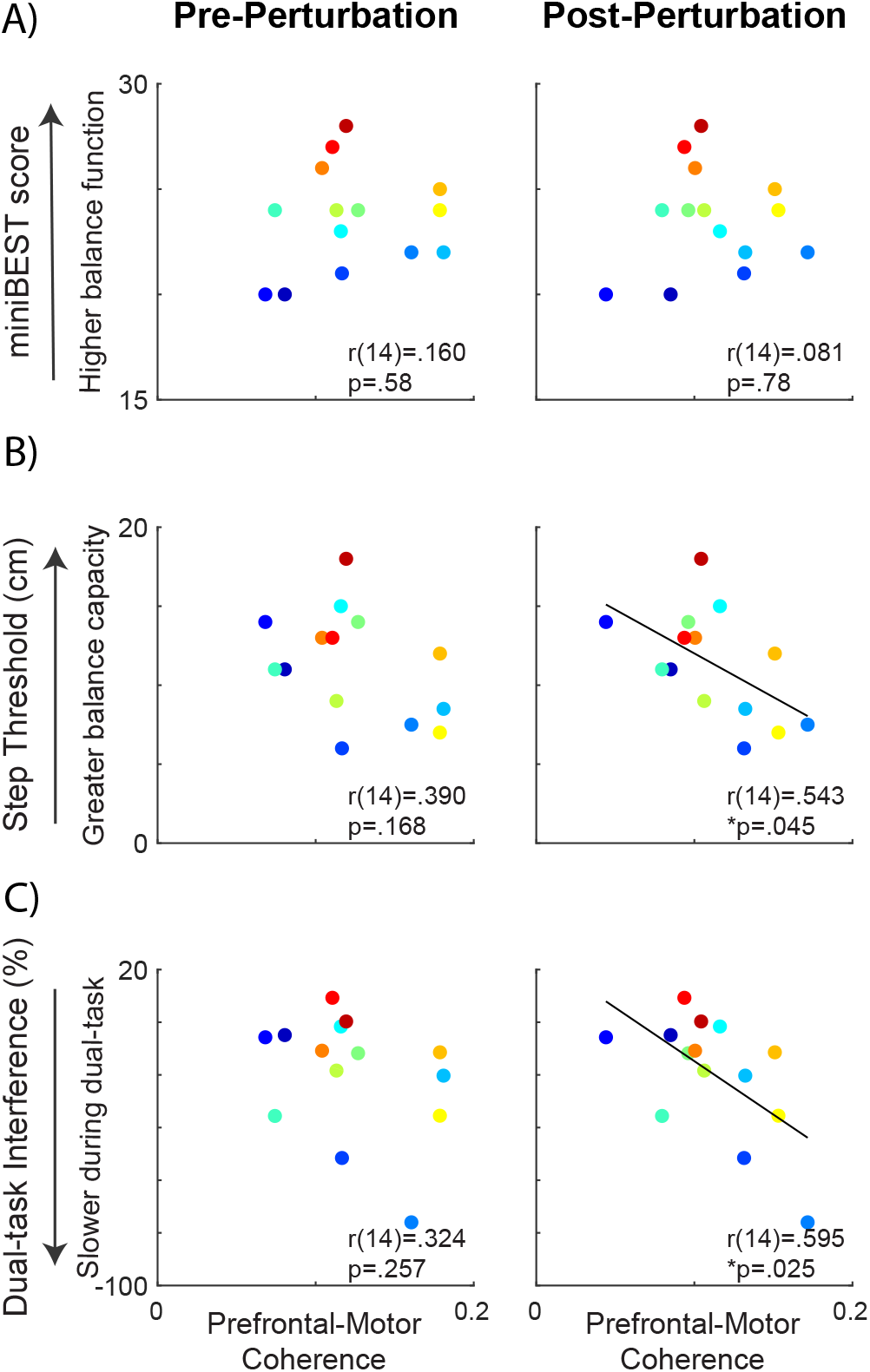
Prefrontal-motor cortical beta coherence pre-and post-perturbation and relationships to balance behavior. Prefrontal-motor coherence was not associated with miniBEST score at any time point (**A**). Prefrontal-motor coherence was negatively associated with reactive step threshold (*p*=.045) (**B**) and cognitive dual-task interference (*p*=.014) (**C**) at post-perturbation but not pre-perturbation time-points.

## Discussion

Our results provide an individualized framework for understanding cortical contributions to balance control, suggesting circuit-specific compensatory roles of cortical engagement in balance control. A key novel finding of the present study is that greater cortical engagement during balance reactions was present in older adults with higher general clinical balance ability, and that interactions between prefrontal and somatosensory regions with motor cortical regions were associated with distinct aspects of balance behavior. Building upon previous research identifying age-related differences in activity and connectivity patterns of these cortical regions, our findings move towards elucidating the role of individual-specific somatosensory-motor and prefrontal-motor cortical circuit engagement for balance control in older adults. Our results yield three main findings: 1) Similar to our previous findings in younger adults (Ghosn et al., 2020), perturbation-evoked beta oscillatory activity over central midline motor cortical regions was negatively correlated with balance ability in older adults, suggesting that perturbation-evoked cortical beta activity may provide a biomarker of individual level of balance challenge across the lifespan; 2) Somatosensory-motor cortical network connectivity during pre-perturbation static standing was positively associated with general clinical balance function, suggesting that somatosensory-motor cortical circuits may play a key role in ongoing sensorimotor integration during standing balance control; and 3) Greater perturbation-evoked prefrontal-motor cortical network connectivity was associated with greater decline in balance performance during cognitive loading and lower threshold for eliciting stepping reactions, implicating that prefrontal-motor cortical circuits may mediate individual reactive balance capacity in older adults. Together, these findings provide evidence that that individual cortical engagement strategies may influence balance behavior in a context-dependent manner. Further, somatosensory-motor and prefrontal-motor circuits show unique timing of contribution to balance control relative to disturbances in standing postural stability.

### Cortical beta power as a biomarker for individual balance challenge

Our findings implicate perturbation-evoked motor cortical beta power as a biomarker for individual balance challenge in older adults. We found that older adults with greater perturbation-evoked beta power had lower general clinical balance ability, measured with the miniBEST (**Figure 3 & 4**). These results are consistent with our previous study in younger adults, who underwent perturbations of larger magnitudes (Ghosn et al., 2020). We specifically observed this relationship in evoked cortical beta power during the later-phase (300-500ms) of the reactive balance response (**Figure 4C**), consistent with the timing of potential later-phase cortical contributions to reactive balance control (B. E. Maki & McIlroy, 2007; Rankin et al., 2000) and also in agreement with younger adults in our previous study (Ghosn et al., 2020). Thus, increased recruitment of cortical resources as level of balance difficulty increases appears to be a strategy utilized by neurotypical younger adults, even in the absence of balance impairment on standard clinical tests (Ghosn et al., 2020). Together with findings of the present study, our results suggest that cortical beta power may reflect a global upregulation of cortical engagement when balance is challenged within an individual across the lifespan. Determination of individual level of balance challenge has important implications for precision medicine approaches to balance treatments and interventions. Motor task practice at precise levels of challenge is a necessary condition to maximize functional cortical neuroplasticity, particularly in older adult populations where, in contrast to young individuals, task practice at the highest levels of motor task challenge can impede skill retention (Bootsma et al., 2020). Thus, the utilization of motor cortical beta activity as a biomarker of individual level of balance challenge (Guadagnoli & Lee, 2004) could serve as a useful clinical tool to tailor individualized treatments and improve balance rehabilitation outcomes for fall prevention.

### Somatosensory-motor network connectivity during standing balance is linked to clinical balance ability in older adults

Stronger functional connectivity of somatosensory-motor circuits prior to perturbation may mediate better overall balance ability, assessed by the miniBEST. In contrast to beta power which appears to reflect overall upregulation of cortical excitability (Aono et al., 2013; Takemi et al., 2013a, 2013b), beta coherence is thought to reflect interactive coupling between two distinct neural sources (Nolte et al., 2004) necessary for information integration and processing (Fries, 2005, 2015; Nolte et al., 2004). We found that older adults who had greater somatosensory-motor coherence during static standing balance prior to perturbation onset had higher miniBEST scores, suggesting that somatosensory-motor cortical circuits may provide an effective mechanism for ongoing sensorimotor processing and integration for balance control in older adults. Interestingly, the relationship between pre-perturbation somatosensory-motor coherence and miniBEST (**Figure 6A**) was the inverse of the relationship between postperturbation motor cortical beta power and miniBEST (**Figure 4C**). Thus, beta power and beta coherence appear to reflect different cortical mechanisms contributing to different aspects of balance control. Somatosensory-motor circuits may have an ongoing influence during standing posture, setting the central nervous system in advance for general balance control and fall avoidance. When greater cortical processing is required to respond to a destabilizing balance perturbation, motor cortical beta power may signal the demand for increased general cortical recruitment.

An upregulation of somatosensory-motor circuit connectivity during standing posture may counteract age-related impairments in peripheral somatosensory system function, as previously postulated (Clark, 2015; Lenz et al., 2012; Pleger et al., 2016). Pervasive declines in sensorimotor processing with aging have been well-documented (Zhang et al., 2011) and may impair the function of fast-acting subcortically-mediated circuits for postural control (Baudry & Duchateau, 2012; Henry & Baudry, 2019; Hortobágyi et al., 2018), contributing to balance dysfunction in older adults (Henry & Baudry, 2019; Hortobágyi et al., 2018). Greater somatosensory-motor coherence in older adults with higher miniBEST scores supports the compensatory role of somatosensory-motor circuit processing for balance control with aging, and is consistent with previous research showing that older adults’ reduced cortical activity, measured with functional near-infrared spectroscopy (fNIRS), during walking when sensory feedback was augmented by wearing textured shoe insoles (Clark, Christou, et al., 2014). Our findings are also in agreement with Malcolm et al. (2020), who found that, in contrast to their younger counterparts, older adults modulated cortical beta activity during quiet static standing as postural difficulty increased with narrowing base of support (Malcolm et al., 2020), further suggesting this cortical strategy may be unique to this age group. Our results build upon this finding, where we delineate a sensorimotor cortical network contribution to this age-related neural compensatory strategy for balance control, potentially identifying specific cortical substrates that could be targeted by precision-medicine strategies to maximize balance function with aging.

### Prefrontal-motor cortical network engagement mediates cognitive interference and reactive balance capacity in older adults

Prefrontal-motor circuit engagement may mediate cognitive dual-task interference and reactive balance capacity in older adults, as we found that individuals with greater perturbation-evoked prefrontal-motor cortical coherence displayed greater slowing of dynamic balance performance during cognitive loading and took reactive steps at lower perturbation magnitudes (**Figure 7**). Our findings compliment those of previous studies suggesting that over-recruitment of prefrontal cortical resources at low levels of motor difficulty imposes a ceiling effect on walking capacity in older adults when walking difficulty is increased with obstacle challenges, particularly after stroke (Clark, Rose, et al., 2014) (Hawkins et al., 2018). Our findings suggest the presence of a similar ceiling effect for prefrontal-motor network engagement in older adults during balance reactions at low levels of perturbation difficulty matched across participants. Our results further expand on these previous findings by identifying the motor circuit specificity of prefrontal networks and applying these circuit-specific mechanisms the context of balance-correcting behavior. Here, older adults with greater interactions between prefrontal and motor cortical regions at a given low magnitude perturbation may have been closer to their “ceiling” for prefrontal-motor cortical engagement. Thus, these individuals may have had reduced availability of cortical resources as balance perturbation magnitude was increased during the reactive step threshold assessment, ultimately necessitating a reactive stepping response at lower perturbation magnitudes. These findings identify prefrontal-motor cortical networks as a potential target for fall prevention strategies aimed specifically at raising the upper-end of individual reactive balance capacity and improving balance performance under cognitive loading conditions.

In the face of a destabilizing postural event, some older adults may capitalize on prefrontal-motor circuit engagement to maintain standing balance, a neural strategy which has distinct behavioral consequences. Perturbation-evoked prefrontal-motor network interactions may reflect executive function and working memory (Naghavi & Nyberg, 2005) and potentially interact as part of a cognitive predictive coding framework (Moran et al., 2014). This prefrontal-motor network engagement during balance reactions may effectively contribute to predictive performance in the aging brain, as it becomes increasingly more accurate in generating predictive models of the environment (Moran et al., 2014), possibly compensating for age-related declines in sensory processing and feedback mechanisms. Another possibility is that recruitment of prefrontal-motor circuits reflects greater focus of attention directed towards balance control when balance is challenged (Boisgontier & Nougier, 2013), though unlike motor cortical beta power, it was not associated with general clinical balance ability measured in the miniBEST. In any case, the engagement of prefrontal-motor circuits during balance reactions may provide some older adults with an effective mechanism to control posture at lower perturbation difficulty levels, enabling them to achieve similar miniBEST scores to individuals who engage different neural control strategies. However, possibly as a result, older adults who rely on prefrontal-motor circuits for balance control tended to show more compromised balance performance when a cognitive demand was placed on the limited pool of executive resources during cognitive dualtask performance (**Figure 5B & 7C**). Our results implicate clinical balance testing under cognitive dual-task conditions as an important adjunct in a battery of clinical balance assessments, as it elucidates the neural strategy that individuals use to support balance control and may predict balance performance under specific task conditions where attentional and executive control resources are concurrently loaded. Further, these findings suggest that rehabilitation interventions for fall prevention that yield little change in clinical miniBEST scores may not necessarily be ineffective. The induced clinical benefits of rehabilitation may come to light in improvements in balance safety, community function, and independence associated with concurrent mobility and cognitive function.

### Role of cortical inhibitory processes in balance control with aging

Cortical beta coherence during balance reactions in older adults may reveal important information about the role of age-related cortical inhibitory processes (Peiffer et al., 2007; Zwergal et al., 2012). Movement-related beta oscillations have been associated with inhibitory GABAergic network activity (Baker, 2007; Muthukumaraswamy et al., 2013), which plays a central role in the modulation of somatosensory processing (Fioravanti et al., 2019). GABAergic networks can precisely influence corticomotor output through synaptic connections with pyramidal neurons, fine-tuning motor control processes (Yamawaki et al., 2008). Over the course of aging, there is a general decline in neural inhibitory processes (Heise et al., 2013; Papegaaij et al., 2014). Heise et al (2013) found that lower intracortical inhibition within the motor cortex during a manual task was correlated with worse motor performance in older adults (Heise et al., 2013). Consistent with these findings, we found that older adults with lower somatosensory-motor beta coherence during standing balance had lower miniBEST scores (**Figure 5A&6A**), suggesting that the preservation of short-range somatosensory-motor inhibitory network connectivity may also contribute to sensorimotor processing and integration mechanisms for standing postural control in older adults.

In the present study, the destabilization of standing balance could have demanded the engagement of inhibitory mechanisms involving the prefrontal cortex for motor suppression of unintentional stepping responses. The prefrontal cortex is a foundational node for the fronto-subthalamic-motor cortical inhibitory network that mediates response-conflict in the motor system (Wessel et al., 2019). If prefrontal-motor beta coherence reflects inhibitory mechanisms associated with this network, individuals with greater interactions between prefrontal-motor regions during balance reactions may have more effectively suppressed stepping reactions in the present study, ultimately yielding higher reactive step thresholds in these older adults. As such, reactive step thresholds may provide a useful clinical probe for inhibitory cortical network function in older adults. Using transcranial magnetic stimulation to directly perturb ongoing motor cortical activity, our lab previously found that event-related motor cortical beta coherence appears to reflect the ability to flexibly modulate cortical network interactions between states of rest and active lower limb muscle contraction; this motor cortical flexibility was linked to functional walking speed capacity that was severely compromised in some older adults after stroke (Palmer et al., 2020). While somatosensory-motor circuits may contribute to baseline standing postural control, our results suggest that the ability of the prefrontal-motor cortical circuits to flexibly engage in balance reactions during a destabilizing postural event may be crucial for upper-end reactive balance control. Here, lower perturbation-evoked prefrontal-motor cortical connectivity in older adults with lower stepping thresholds could stem from an age-related decline in inhibitory network function and consequently reduce the flexible modulatory capacity of the motor system to respond to destabilizing balance events. It remains unclear why some individuals may be more resistant to age-related declines in neural inhibitory processes. More research is needed to improve our understanding of the underlying causal factors for differences in cortical circuit engagement during reactive balance between individuals in aging populations. However lifestyle behaviors, particularly physical activity, appear to play a key role in the slowing of normal age-related declines in inhibitory neural network function (Levin & Netz, 2015).

### Limitations

The present study utilized a channel-based beta power and coherence approach as a first step to quantify functional connectivity during balance reactions and enhance the clinical translation of neural mechanistic findings. However, this approach limits the spatial anatomical specificity of EEG analyses, raising the possibility that findings of the present study could be influenced by subcortical and cortical regions outside of the somatosensory, motor, and prefrontal cortical regions. Given the close proximity of the channels overlying these cortical regions, we employed a conservative IPC approach for functional connectivity analyses that excludes instantaneous common source activity. Though this approach increases the confidence that coherence measures reflect a greater degree of true neural interactions between sources, a limitation of this analysis is that we cannot determine directionality of these neural interactions. It is likely that somatosensory-motor and prefrontal-motor coherence measures here reflect both feedforward (somatosensory-to-motor and prefrontal-to-motor) and feedback (motor-to-somatosensory and motor-to-prefrontal) network interactions. Future studies could assess directionality of these interactions with effective connectivity approaches. During cognitive dual task performance assessment, we quantified motor behavioral performance, but cognitive performance was not quantified. Future studies may address this limitation by quantifying concurrent declines in balance and cognitive performance that have been reported in older adults.

## Conclusions

Our findings show that older adults engage individual circuit-specific cortical strategies during balance behavior that are linked to distinct aspects of balance control. Whereas both somatosensory-motor and prefrontal-motor circuits can contribute to balance function across individuals, prefrontally-mediated cortical strategies may be less effective for balance control in distracting contexts and under the most challenging balance conditions. These cortical strategies may each have unique contributions to balance behavior depending on the context (e.g., concurrent cognitive load) and/or the environment (e.g., level of balance difficulty). Our findings potentially identify suboptimal neural control strategies that could be targeted for early intervention to prevent emergence of clinical balance deficits in aging populations.

## Acknowledgements and Funding

We would like to acknowledge Kamal Shadi for assisting in data analysis. This work was supported by postdoctoral fellowship awards from the American Heart Association American Heart Association [AHA00035638] and the National Institutes of Health [F32HD096816] to JAP, grants from the National Institutes of Health [K12HD055931] to MRB, [5T90DA032466, 1P50NS098685, R01 HD46922] to LHT, and a grant from the National Science Foundation [1137229] to LHT.

